# CRISPR-Cas9-mediated genome engineering exaggerates genomic deletion at 10q23.31 including the *PTEN* gene locus mimicking cancer profiles

**DOI:** 10.1101/2023.04.05.535680

**Authors:** Keyi Geng, Lara G. Merino, Raül G. Veiga, Christian Sommerauer, Janine Epperlein, Eva K. Brinkman, Claudia Kutter

**Affiliations:** Department of Microbiology, Tumor, and Cell Biology, Karolinska Institute, Science for Life Laboratory, Sweden

**Keywords:** CRISPR-Cas9, PTEN, 10q23.31, genomic deletion, cancer

## Abstract

The CRISPR-Cas9 system is a powerful tool for studying gene functions and has tremendous potential for disease treatment. However, precise genome editing requires thorough assessments to minimize unintended on- and off-target effects. Here, we report an unexpected deletion of a 287 kb region on Chromosome 10 (10q23.31) in chronic myelogenous leukemia HAP1 cells, which are frequently used in CRISPR screens. The deleted region encodes regulatory genes, including *PAPSS2, ATAD1, KLLN*, and *PTEN*. We found that this deletion was not a direct consequence of CRISPR-Cas9 off-targeting but rather occurred frequently by the process of generating CRISPR-Cas9-modifed cells. The deletion was associated with global changes in histone acetylation and gene expression, affecting fundamental cellular processes such as cell cycle and DNA replication. We detected this deletion in cancer patient genomes. As in HAP1 cells, the deletion contributed to similar gene expression patterns among cancer patients despite interindividual differences. Overall, our findings suggest that the unintended deletion of 10q23.31 can confound CRISPR-Cas9 studies, highlights the importance of assessing unintended genomic changes in CRISPR-Cas9-modified cells and may have clinical significance in cancer research.

**Highlights:**

- CRISPR-Cas9-modified HAP1 cells carry an unexpected large genomic deletion at 10q23.31 encompassing four protein-coding genes frequently expressed across various cell types.
- The 10q23.31 deletion is accompanied by global changes in histone modification and transcriptomes.
- The generation of CRISPR-Cas9-modified cells rather than Cas9 activity increases the frequencies of the deletion at 10q23.31.
- The 10q23.31 deletion identified in HAP1 cells resembles a commonly occurring deletion pattern in cancer patients.

## INTRODUCTION

The clustered regularly interspaced short palindromic repeats (CRISPR) and CRISPR-associated (Cas) system is a widely used genome engineering technology because of its simple programmability, versatile scalability, and targeting efficiency (1). Although researchers are rapidly developing CRISPR-Cas9 tools, the biggest challenge remains to overcome undesired on- and off-targeting outcomes. Previous studies reported undesired genomic alterations resulting in the integration of functional target-derived sequences (2), large deletion at the double-strand break (DSB) site (3), chromothripsis (4), segmental chromosomal losses (5) or translocations (6) that often co-occur (2). Most of these genomic rearrangements remain undetectable by conventional validation methods, which emphasizes the need to thoroughly assess CRISPR-Cas9-modified cells or organisms by more advanced tools.

The complexity of genomic outcomes is linked to the experimental model system. For example, dysfunctional repair mechanisms in certain cell lines can influence the cellular preference for employing a specific repair pathway, which can result in different editing outcomes (7). Furthermore, in polypoid cells, several on-target alterations can occur on multiple alleles resulting in varying phenotypes and biological consequences across cells (2). Therefore, the choice of the cellular model system is crucial for investigating biological processes or diseases.

The chronic myeloid leukemia-derived HAP1 cell line is often used in genetic studies and large-scale CRISPR screens because its near-haploid genotype increases the probabilities of acquiring modified cells with homozygous genotypes. Homozygous editing is often preferred when biological functions of the CRISPR-Cas9-modified target are assessed (8, 9). In contrast, a targeted region or gene may only be successfully modified on one allele in diploid or polyploid cells. Thus, the remaining functional copy on the other allele(s) can mask the effect of the CRISPR-Cas9-modified copy, preventing identification of measurable phenotypic changes and gene functionality. Since HAP1 is widely used in CRISPR-Cas9 applications, it is crucial to eliminate any confounding genetic variances that could jeopardize the conclusions drawn from the experiments.

In this work, we report the inconsistent occurrence of a large and unexpected genomic deletion at 10q23.31 in HAP1 cells. This deletion was accompanied by wide-spread changes on the chromatin level and in gene expression. Instead of a commonly reported gRNA-dependent off-targeting event, we found that the generation of CRISPR-Cas9 edited cells and/or exposure to cellular stressors greatly increases the occurrence of the deletion. We found the 10q23.31 deletion in HAP1 cells but also detected its frequent occurrence across several cancer types in patients. Our findings highlight the importance to consider collateral deletions when assessing mechanistic functions of genes or regulatory regions in cells commonly used for basic research or newly isolated from patients for personalized medicine.

## RESULTS

### CRISPR-Cas9-modified HAP1 cells contained an unexpected 10q23.31 deletion

To study two proximal transfer RNA (tRNA) genes on Chromosome (Chr) 17 (17q12), we previously used the dual gRNA system to generate two variants of CRISPR-Cas9-modified and single cell-derived HAP1 clones in which the genomic region with (Δt clones) or in between (Δi clones) two tRNA genes was removed (**Fig. 1A**) (2). We profiled genomic occupancy of histone (H) 3 lysine (K) 4 trimethylation (H3K4me3) and K27 acetylation (H3K27ac) by ChIP-seq. As intended, H3K4me3 and H3K27ac ChIP-seq enrichment and background signals were absent in between the targeted regions on Chr 17 in the Δt1 and Δi17 clones but not in the CRISPR-Cas9-unmodified control (ctrl) clone confirming the successful deletion of the corresponding genomic regions (**Fig. 1A, Supplementary Fig. S1A**). Unexpectedly, ChIP-seq enrichment or background signals were also undetectable on Chr 10 (10q23.31) in the Δt1 and Δi17 clones but not in the unmodified cell clone (**Fig. 1B**). Since the guide RNA sequences (gRNAs) used for generating the Δt and Δi cell clones were different, we concluded that the 10q23.31 deletion occurred in a gRNA-independent manner (**Supplementary Table S1**).

**Fig 1.**
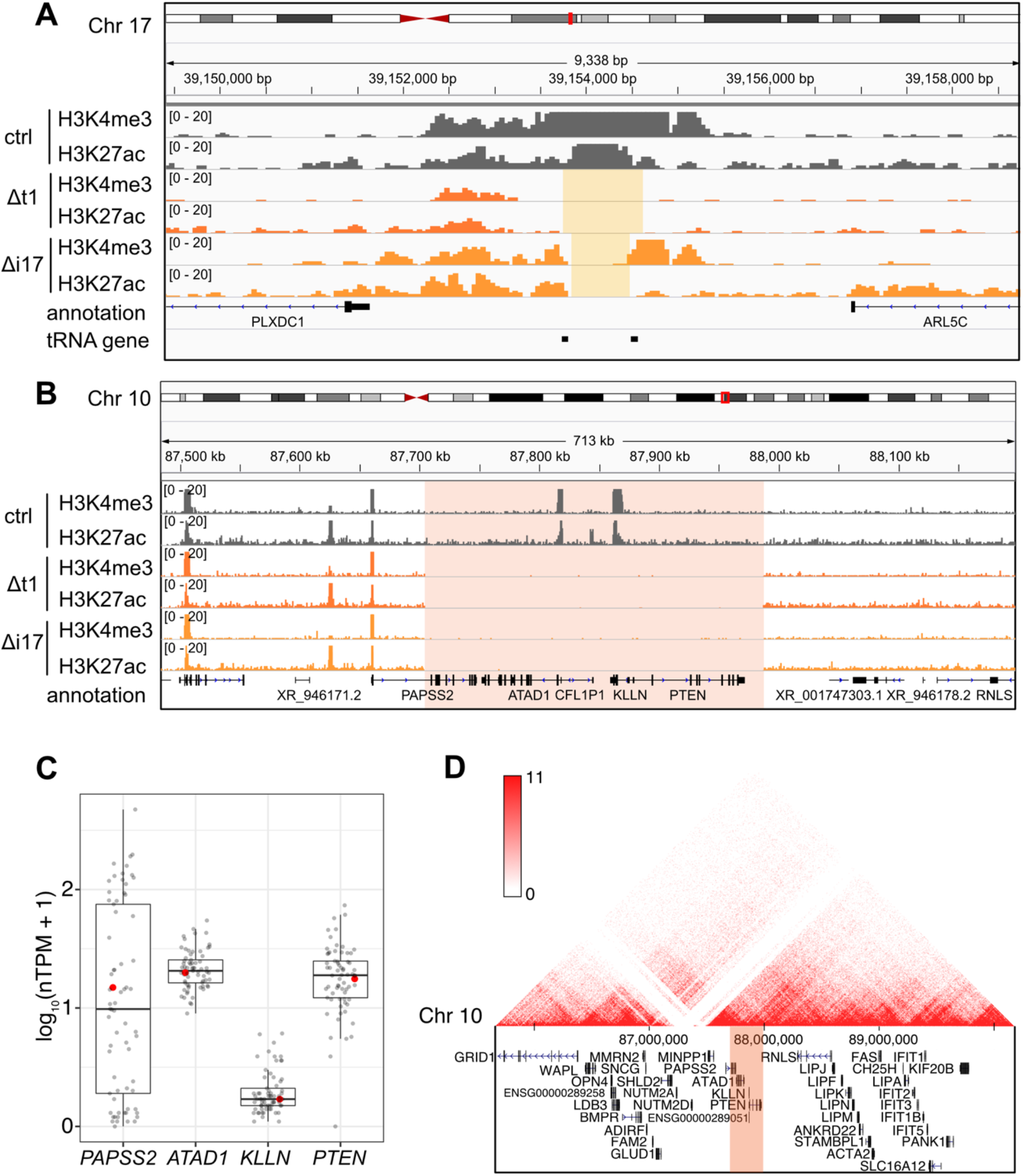
The loss of the *PAPSS2-PTEN* locus occurred in various CRISPR-Cas9-modified genotypes in HAP1 cells. (**A-B**) The hg38 genome browser view shows normalized H3K4me3 and H3K27ac ChIP-seq reads in control (ctrl, dark grey), Δt1 (orange) and Δi17 (yellow) HAP1 cell clones at (**A**) the targeted gene locus on Chr 17 (beige box) and (**B**) the *PAPSS2-PTEN* locus on Chr 10 (red box). (**C**) Box plot shows the normalized gene expression values (in normalized TPM) of the four protein-coding genes within the deleted region on Chr 10. Each dot represents gene expression values across different cell lines (grey) including HAP1 (red). Normalized TPM values were obtained from the Human Protein Atlas (v. 21.0). Median (horizontal line), interquartile range (box), lower and upper quartile (whiskers) are shown. (**D**) The genome browser presents a 3D-contact map of Hi-C data at the *PAPSS2-PTEN* locus in HAP1 cells (genomic contact frequency, red: high, white: low). The deleted region is highlighted (red box).

To further assess the frequency of the 10q23.31 deletion, we inspected our other CRISPR-Cas9-modified HAP1 cell clones by PCR using primers annealing within the deleted genomic region (**Supplementary Table S1**). The absence of a PCR amplicon in 61% (14/23) Δt and 44% (4/9) Δi clones suggested frequent losses of the 10q23.31 region in our CRISPR-Cas9-modified cell clones (**Supplementary Fig. S1B**).

The deleted 10q23.31 locus of about 287 kb encompassed four protein-coding genes (*PAPSS2, ATAD1, KLLN* and *PTEN*) and one pseudogene (*CFL1P1*). The four protein-coding genes were widely expressed in 69 cell lines of different tissue origin (**Fig. 1C**) and exert essential molecular functions. As previously reported, the PAPSS2 enzyme controls the sulphate activation pathway (34, 35), the transmembrane helix translocase ATAD1 removes mislocalized proteins from the mitochondrial outer membrane (36), KLLN regulates cell cycle and apoptosis (37), and the tumor suppressor PTEN acts as a protein phosphatase (38, 39). Furthermore, Hi-C and ChIA-PET data obtained from HAP1 and other cell types revealed multiple long-range interactions between the 10q23.31 and other genomic regions indicating additional roles in three-dimensional (3D) gene regulation (**Fig. 1D, Supplementary Fig. S1C**). Lastly, according to our H3K4me3 and H3K27ac profiling, the deletion ranged from the first intron of the *PAPSS2* to a genomic site downstream of the *PTEN* gene. We therefore referred to this genomic deletion as Δ*PAPSS2-PTEN*.

In sum, the *PAPSS2-PTEN* gene locus was frequently deleted in CRISPR-Cas9-modified cell clones. Given that this region is important in 3D genome organization and encodes four protein-coding genes with essential molecular functions, its unintended deletion may dominate over the expected gRNA-mediated CRISPR-Cas9 genomic deletion and could lead to biological misinterpretations.

### *ΔPAPSS2-PTEN* cells showed abnormal transcript signatures

Since we identified the deletion of the *PAPSS2-PTEN* locus through chromatin profiling, we next examined the impact of the deletion on the transcriptome. In alignment with our ChIP-seq results, our RNA-seq data confirmed the complete loss of gene expression at the *PAPSS2-PTEN* locus in the Δt1 clone when compared to the control cell clone (**Fig. 2A, top 4 tracks**). Further inspection revealed reads mapping to the first exon of *PAPSS2* confirming that the genomic region including the promoter, transcriptional start site (TSS) and first exon of *PAPSS2* remained intact (**Fig. 1B, Fig. 2B, top 2 tracks**). Furthermore, we found reads mapping to the positive strand downstream of the *PTEN* gene body in the Δt1 clone but not in the control cell clone (**Fig. 2C, top 2 tracks**). This transcript signature was likely caused by polymerase II (Pol II) readthrough from the altered *PAPSS2* gene locus since Pol II can still be recruited to the *PAPSS2* promoter and initiated aberrant transcript formation in Δ*PAPSS2*-*PTEN* cells. However, the *PAPSS2* Pol II termination signal as well as major parts of the *PAPSS2* gene body were lost together with *ATAD1, KLLN* and *PTEN*.

**Fig 2.**
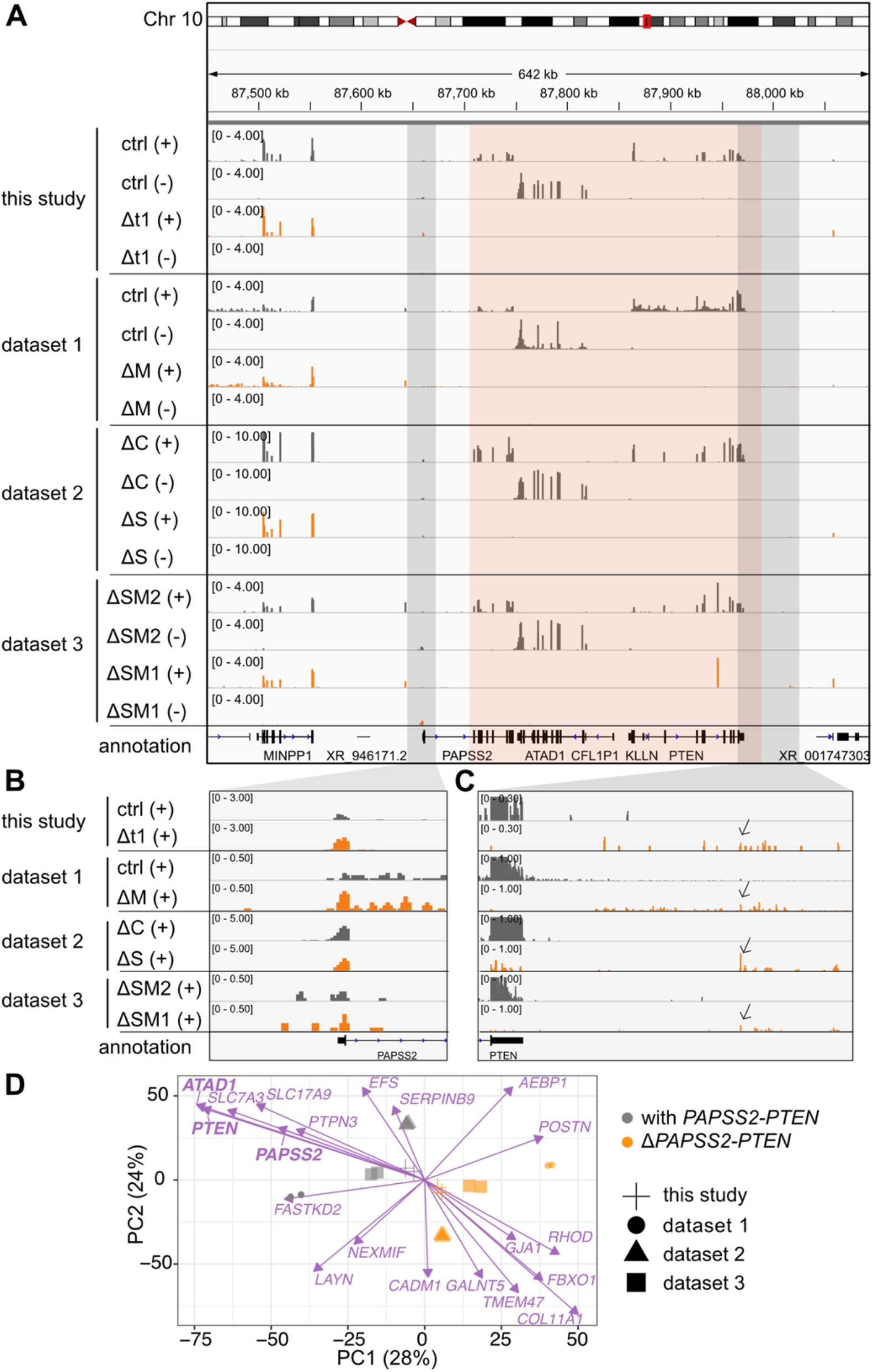
The unintended loss of the *PAPSS2-PTEN* locus resulted in similar transcriptional changes despite genotypical differences in CRISPR-Cas9-modified HAP1 cell clones. (**A-C**) The hg38 genome browser view shows normalized RNA-seq coverage tracks for the plus (+) and minus (-) strand over the *PAPSS2-PTEN* locus (highlight in red box) for CRISPR-Cas9-modified (dark grey) and control (ctrl) HAP1 cell clones (orange) generated in this and other studies (dataset 1 to 3). (**B-C**) The (**B**) upstream and (**C**) downstream region of the *PAPSS2-PTEN* locus (indicated as light grey boxes in Fig. 2A) are magnified and visualized for the plus strand. Pol II read-through signals are indicated (arrows). (**D**) Factorial map of the PCA after batch effect correction is shown for the top 20 genes (purple arrows) separating the HAP1 cell clones that contain (with, grey circle) or lost (Δ, orange circle) the *PAPSS2-PTEN* locus in our and other datasets (geometric shapes) in PC1 and PC2. The proportion of variance explained by each PC is indicated in parenthesis.

To confirm our newly identified genome and transcript signature at the Δ*PAPSS2-PTEN* region, we searched for published RNA-seq data of CRISPR-Cas9-modified HAP1 cells. We retrieved RNA-seq datasets from three independent studies in which various genes were modified, affecting the *METAP1* (ΔM) (dataset 1), *C12orf49* (ΔC) and *SREBF2* (ΔS) (dataset 2) as well as *SMARCC1* (ΔSM1) and *SMARCC2* (ΔSM2) (dataset 3) genes (**Supplementary Table S2**). No direct network interactions were found between those CRISPR-Cas9-modified genes and the Δ*PAPSS2-PTEN* encoded genes (**Supplementary Fig. S2A**). Similar to our Δt1 clone, we also found that the *PAPSS2-PTEN* locus was deleted in the ΔM, ΔS and ΔSM1 but not in the ΔC and ΔSM2 clones (**Fig. 2A-B**).

To identify common gene expression signatures associated with the deletion of the *PAPSS2-PTEN* locus, we combined our and the other three datasets. We performed a principal component analysis (PCA). As commonly observed, PC1 and PC2 separated the samples by dataset likely due to technical biases (40), such as sample and library preparation or sequencing (**Supplementary Fig. S2B**). However, the subsequent PCs (PC3 to PC5) showed that samples with the *PAPSS2-PTEN* locus deletion (Δ*PAPSS2-PTEN* group) clustered despite their genotypic differences caused upon the deliberate CRISPR-Cas9 gene modification (**Supplementary Fig. S2C-D**). We corrected for batch biases to minimize the differences between the datasets that were introduced during the sample and library preparation. After the batch correction, PC1 and PC2 grouped the samples in either the *PAPSS2-PTEN* deletion positive (Δ*PAPSS2-PTEN*) or negative (with *PAPSS2-PTEN*) group (**Fig. 2D**). We next examined the top 20 genes contributing to PC1 and PC2. *ATAD1, PTEN* and *PAPSS2* were among the top genes that contributed to the strongest separation of the two groups (**Fig. 2D**). None of the top genes included the originally indented gene knockouts (**Fig. 2D**) and were located on different chromosomes (**Supplementary Table S4-5**).

Thus, our inspection of transcriptome data from various CRISPR-Cas9-modified HAP1 cell clones confirmed a gRNA-independent deletion at the *PAPSS2-PTEN* locus. Importantly, the unexpected *PAPSS2-PTEN* deletion resulted in similar gene expression changes dominating over the intended gene modification, which could bias the assessment of gene functionality.

### Gene expression changes in Δ*PAPSS2-PTEN* HAP1 cells affected fundamental processes including cell cycle and DNA replication

To obtain a comprehensive understanding of the transcriptional changes in Δ*PAPSS2-PTEN* HAP1 cells, we performed a differential gene expression analysis. We found a total of 2,918 differentially expressed (DE) genes corresponding to 1,489 down- and 1,429 upregulated genes (**Fig. 3A**) located on different chromosomes with no apparent positional clustering (**Fig. 3B**). A few DE genes were located on Chr 10 but resided in topologically associating domains (TADs) other than the TAD encompassing the *PAPSS2-PTEN* locus. For example, *SNCG* was located in linear distance closest (700 kb) and *MALRD1* furthest (6,800 kb) to the *PAPSS2-PTEN* locus (**Fig. 3C, Fig. 1D**). These results suggested that complex transcriptional changes observed in Δ*PAPSS2-PTEN* cells were caused through regulatory networks rather than local impacts of the deletion on nearby gene expression.

**Fig 3.**
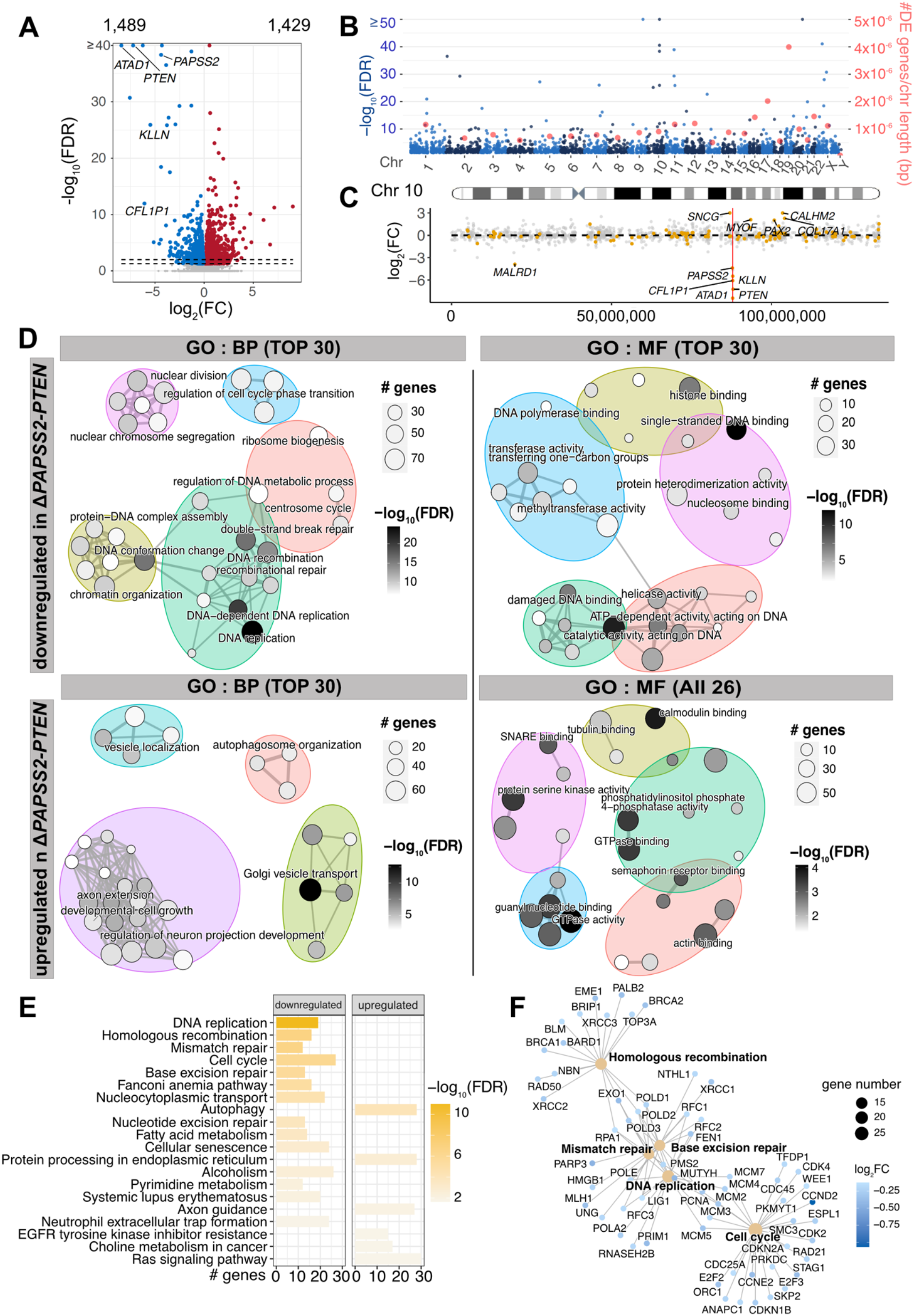
The *PAPSS2-PTEN* locus deletion was associated with confounding transcriptomic alterations impacting molecular processes. (**A**) Volcano plot separates DE genes when comparing transcriptomes of HAP1 cell clones with and without the *PAPSS2-PTEN* deletion. Each dot shows non-DE genes (grey) or DE genes (red: log_2_(fold-change, FC)>0 and blue:log_2_(FC)<0, FDR-adjusted *p* values, FDR≤0.05). The numbers of up- and downregulated genes are labelled above the plot and genes located in the deleted 10q23.31 region are highlighted. The two dashed lines indicate FDR-adjusted *p* values (FDR=0.05 and FDR=0.01). (**B**) Manhattan plot shows the chromosomal distribution (light and dark blue dots) and frequency of the DE genes normalized by chromosome length (red dots). (**C**) Dot plot shows the distribution on Chr 10 (x-axis) and fold-change (y-axis) of non-DE (grey) and DE genes (orange). The red line highlights the location of the deleted *PAPSS2-PTEN* locus. DE genes with |log_2_(FC)| >2 are labelled. (**D**) Enrichment maps illustrate the up to top 30 or 26 significantly enriched biological process (BP) and molecular function (MF) GO terms for down- and upregulated DE genes. Each node represents one enriched GO term. The sizes of the nodes are determined by the numbers of DE genes contributing to the enriched GO term. The nodes are colored based on FDR-adjusted *p* values. (**E**) Bar plot shows the top 20 enriched KEGG pathways for down- and upregulated genes (ranked by FDR-adjusted *p* values). (**F**) The gene concept network displays genes contributing to the top 5 enriched KEGG pathways (ranked by FDR-adjusted *p* values). The node diameter determines the number of genes, colored according to the fold-change.

To assess whether specific regulatory processes were changed upon the deletion of the *PAPSS2-PTEN* locus, we grouped DE genes according to gene ontology (GO) terms (**Fig. 3D, Supplementary Table S6-9**). The downregulated genes were significantly enriched for biological processes, such as DNA replication, cell cycle and DSB repair as well as molecular functions, including DNA- and histone binding. In contrast, the upregulated genes significantly controlled biological processes and molecular functions linked to GO terms such as development and catalytic activities. Accordingly, our KEGG pathway enrichment analysis confirmed the identified GO terms (**Fig. 3E, Supplementary Table S10-11**). Many DE genes, such as replication factor C (*RFC1, RFC3* and *RFC3*), the *MCM* gene family as well as genes forming DNA polymerase subunits, controlled crucial cell processes, and contributed to the top 5 enriched pathways (**Fig. 3F**).

Since our batch effect correction could have introduced biases, we performed the DE analysis for each dataset separately by comparing the groups with and without the *PAPSS2-PTEN* locus. The large majority (88%, 2,570 of 2,918) of DE genes was commonly deregulated between the pooled batch effect corrected datasets and in each separately analyzed dataset (**Supplementary Fig. S2E**). Importantly, DE genes identified in at least three of the four separately analyzed datasets were significantly enriched in biological processes, comprising cell cycle, DNA replication, DNA repair and molecular functions, including single-strand DNA binding, which were in accordance with the results obtained after batch effect correction (**Supplementary Fig. S2F**).

In conclusion, transcriptome signatures in CRISPR-Cas9-modified cell clones carrying the genomic deletion of the *PAPSS2-PTEN* locus were profoundly altered.

### Gene expression and H3K27ac changes were linked in Δ*PAPSS2-PTEN* HAP1 cells

Since our GO enrichment analysis suggested changes in chromatin organization and modification, we expected alterations in genome accessibility in the *ΔPAPSS2-PTEN* cell clones. We therefore quantified genomic occurrences of H3K27ac, which demarcates active promoters and enhancers (41). We used H3K27ac data obtained from our HAP1 Δt1 (Δ*PAPSS2-PTEN*) and control (with *PAPSS2-PTEN*) cell clones (**Fig. 1A-B**). In addition, we retrieved publicly available H3K27ac data from the HAP1 cell clones ΔSM1 (Δ*PAPSS2-PTEN*) and used Δ*SMARCC4* (ΔSM4, with *PAPSS2-PTEN*) as control (**Fig. 2A, Fig. S3A, Supplementary Table S3**). After identifying genomic regions enriched for H3K27ac, we performed a PCA that separated the samples first, by data source (PC1 with 68% variance) and second, by the *PAPSS2-PTEN* genotype (PC2 with 11% variance) (**Fig. 4A**).

**Fig 4.**
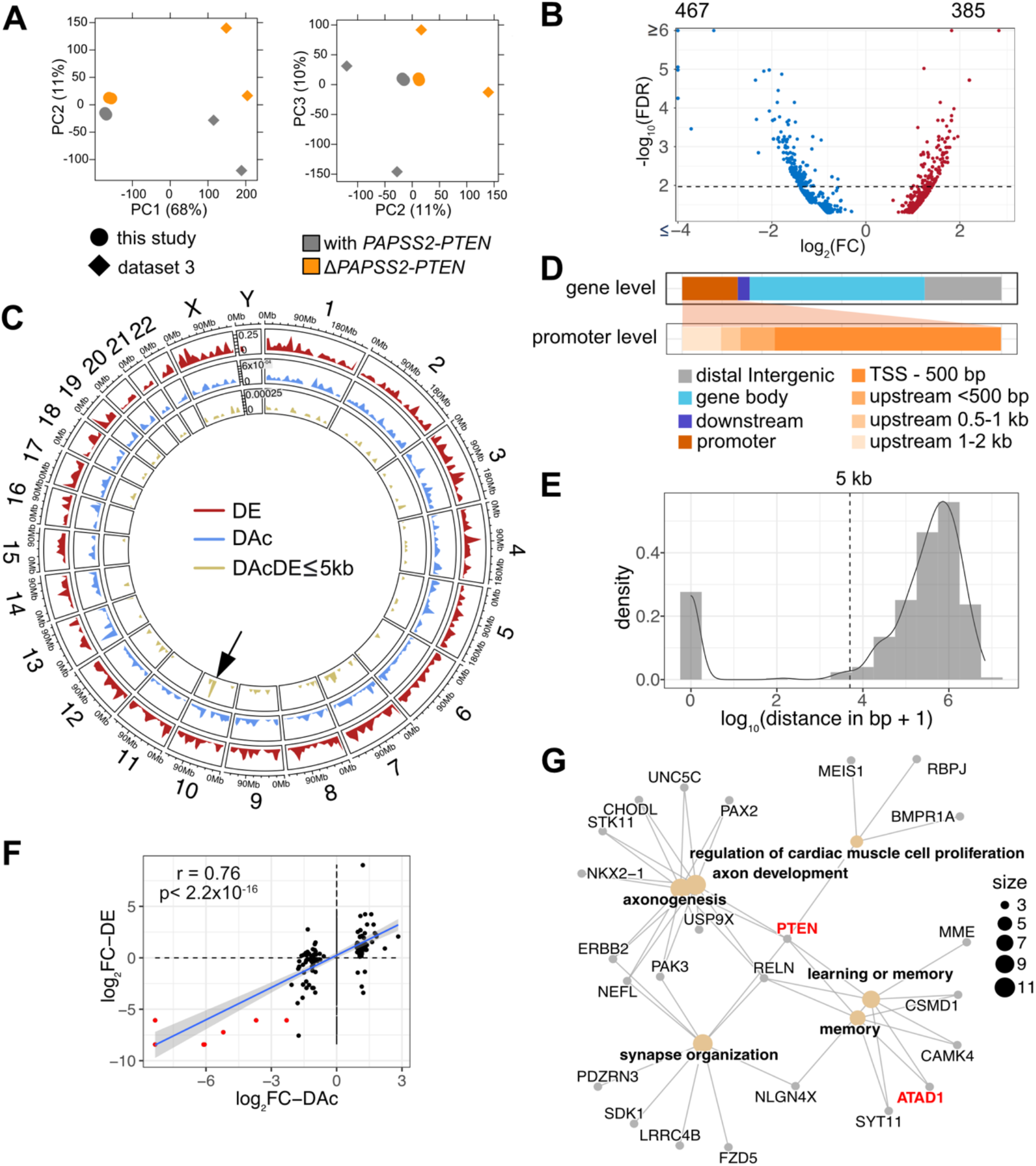
H3K27ac occupancy was altered in the genomes of *PAPSS2-PTEN* deleted HAP1 cells. (**A**) Factorial maps of the PCA of global H3K27ac ChIP-seq signals separates samples according to absence (orange) or presence (grey) of the *PAPSS2-PTEN* locus in HAP1 CRISPR-Cas9 deletion clones generated in this (circle) or other (diamond) studies. The proportion of variance explained by each PC is indicated in parenthesis. (**B**) Volcano plot shows differentially acetylated H3K27ac ChIP-seq peaks (DAc) (dashed line corresponds to FDR=0.01). Number of decreased (blue) and increased (red) acetylated H3K27 peaks are labelled (top). (**C**) Circular representation of the human genome illustrates each chromosome proportionally scaled to its length. Tracks inserted in the circle show the genomic location and frequency of DE genes (red), DAc genomic regions (blue), and DAc peaks with DE genes located nearby (genomic distance DAc to DE, DAcDE ≤ 5 kb, green). Arrow highlights DAcDE pairs on Chr 10. (**D**) Stacked bar plots demonstrate proportional frequencies of DAc peaks for genomic features (color-coded). Subcategories for promoter features are further divided. (**E**) The density plot shows the distance of individual DAc peak to the nearest differentially expressed gene. The dashed line indicates the 5,000 bp cut-off. (**F**) Dot plot correlates fold-changes in DAc peaks and nearby DE genes (distance ≤5 kb). The grey area within the plot represents the 95% confidence interval (linear model). Spearman’s rank correlation coefficient (r) and *p* value (p) are indicated. The DAcDE pairs are colored (red) if located within the deleted 10q23.31 locus. (**G**) The gene concept network shows DE genes with a nearby (≤ 5 kb) DAc peak grouped into significantly enriched GO terms. The size of the node is determined by the number of DE genes contributing to the enriched GO term.

Our subsequent differential enrichment analysis uncovered 852 differentially acetylated (DAc) H3K27 regions, that were distributed on all chromosomes (**Fig. 4B-C, Supplementary Fig. S3B**). About 17% (147/852) of the DAc peaks resided in promoter regions, mostly within 500 bp downstream from the nearest annotated TSSs (**Fig. 4D**). Over half (54%, 463/852) of the DAc peaks were located within gene bodies and nearly a quarter (24%, 204/852) in intergenic regions (**Fig. 4D**). To define the influence of differential H3K27 acetylation on gene expression in Δ*PAPSS2-PTEN* cells, we calculated the distances between each DAc peak and the closest DE gene (**Fig. 4E**). About 85% (730/861) of the DAc peaks were located distant (>5 kb) from any DE genes. We restricted our subsequent analysis to the 131 DAc peaks (15%, 131/861) that were adjacent to or overlapping with the DE genes (≤ 5 kb). These DAcDE pairs were spread across many chromosomes and were enriched on Chr 10 that encompassed the *PAPSS2-PTEN* locus among others (**Fig. 4D**). Differential H3K27 acetylation and gene expression levels of the DAcDE pairs highly correlated (**Fig. 4F**), likely due to DAc of H3K27 in the DE promoters and gene body (**Supplementary Fig. S3C**). DE genes with altered H3K27ac levels were enriched in six GO terms linked to proliferation, development and memory (**Fig. 4G**). *PTEN* connected all and *ATAD1* 33% (2/6) GO terms.

Thus, irrespective of the actual genomic CRISPR-Cas9 modification, we found that the unintended deletion of the *PAPSS2-PTEN* locus in HAP1 cells was related to dramatic changes on the chromatin and transcript level.

### Generation of CRISPR-Cas9 deletion clones exaggerated the loss of the *PAPSS2-PTEN* locus

To explain the cause of the *ΔPAPSS2-PTEN*, we systematically tested individual steps commonly performed when generating CRISPR-Cas9 deletion clones. First, we had already ruled out a gRNA-mediated off-targeting effect since the unintended *PAPSS2-PTEN* deletion was consistently detectable when a variety of different genomic regions were targeted (**Fig. 2**). To empirically exclude that Cas9 cuts the *PAPSS2-PTEN* locus in a gRNA-independent manner, we transfected HAP1 cells with a CRISPR-Cas9 plasmid encoding the puromycin resistance gene used for antibiotic-based clonal selection without (px459) or with gRNA sequences (px459+gRNA_Δt) (**Fig. 5A-B**). We assessed the frequency of the *PAPSS2-PTEN* locus deletion in each single cell-derived clone by PCR or qPCR using primers binding to the *ATAD1* or *PTEN* promoter region (**Fig. 5A, Supplementary Table S1**). We detected insignificant differences in the frequency of the *PAPSS2-PTEN* deletion in HAP1 cell clones transfected without (67%, 12/18) and with (83%,15/18) the gRNA-containing CRISPR-Cas9 plasmid (Fisher’s exact test, *p*=0.443) (**Fig. 5B**). This result verified that the genomic deletion of the *PAPSS2-PTEN* locus was gRNA-independent and likely caused during the generation of CRISPR-Cas9 deletion clones.

**Fig 5.**
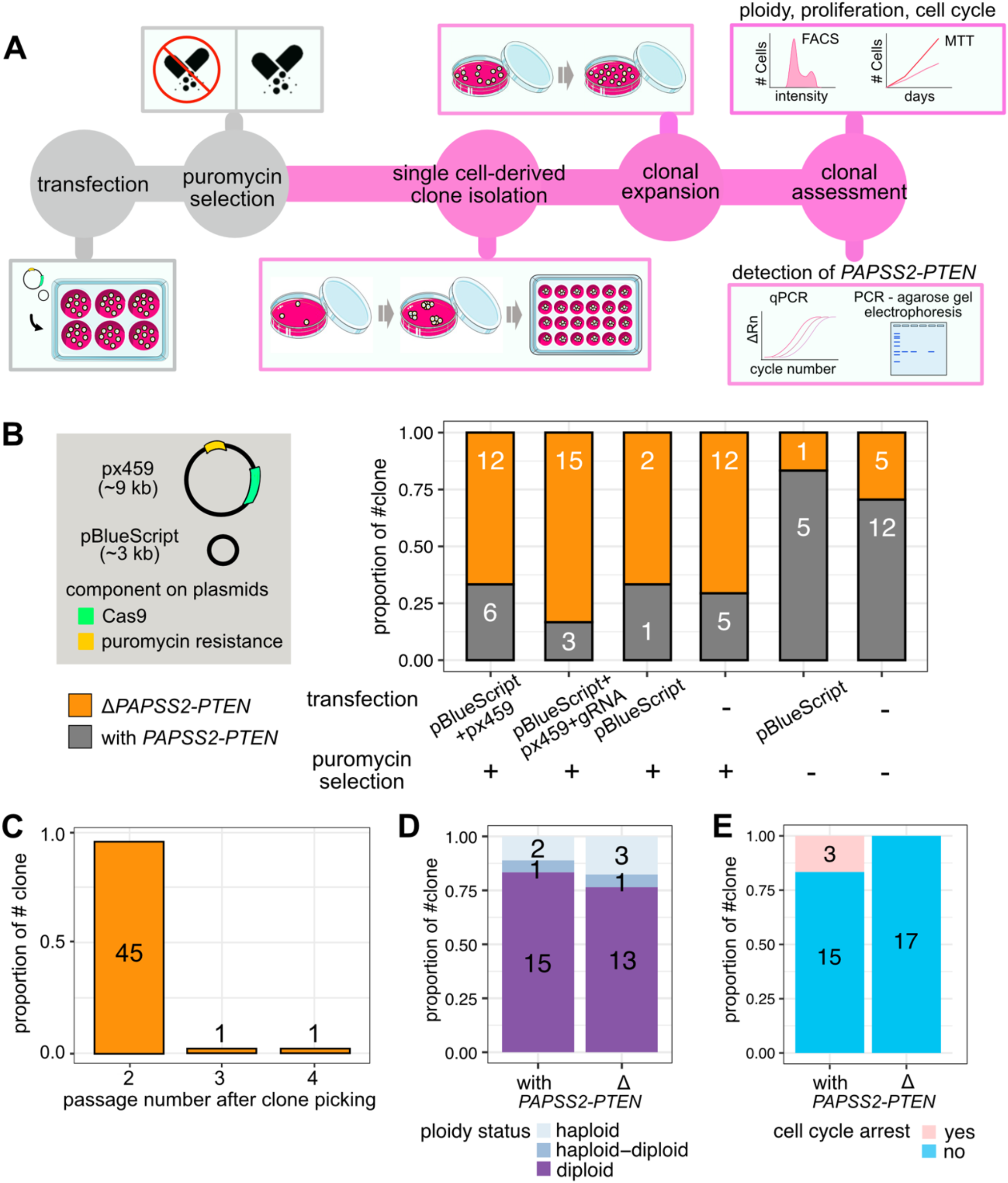
The generation of CRISPR-Cas9-modified clones exacerbated the loss in the 10q23.31 region resulting in cell cycle changes. (**A**) Schematic illustration displays the workflow for determining the frequency of the genomic deletion of the *PAPSS2-PTEN* locus and associated cellular consequences in single cell-derived HAP1 clones. The transfection and puromycin selection steps (colored in grey) were omitted as part of the testing in some experiments. (**B-E**) Bar plots show the frequency of single cell-derived clones without (orange) and with (grey) the *PAPSS2-PTEN* locus (**B**) upon transfection with various plasmids (*left*, green: Cas9 gene, yellow: puromycin resistance gene) with or without puromycin selection, (**C**) over several cell passages after single cell selection, (**D**) in different ploidy stages (light blue: haploid, blue: haploid-diploid, purple: diploid) and (**E**) arrested during cell cycle progression (pink: arrested, blue: not arrested). The number of cell clones for each group is indicated.

Second, since we co-transfected the large-size CRISPR-Cas9 plasmid with a small-size plasmid (pBlueScript) to enhance the transfection efficiency (10), we assessed the occurrence of the *PAPSS2-PTEN* deletion in HAP1 cell clones transfected with or without the small-size plasmid (pBlueScript) that did not encode a puromycin resistance gene. Genomic *PAPSS2-PTEN* locus deletions were detectable in HAP1 cell clones transfected with only the pBlueScript (67%, 2/3) or no plasmid (71%, 12/17) (**Fig. 5B**). This result further supported that the *PAPSS2-PTEN* locus deletion in HAP1 cell clones resulted neither through Cas9 off-targeting nor pBlueScript co-transfection.

Lastly, we omitted the delivery of CRISPR-Cas9 components and the puromycin selection step. Instead, we transfected HAP1 cells with either only the pBlueScript plasmid or no plasmids and propagated the cells in puromycin-free medium. Omitting of the antibiotic selection step reduced the occurrences of the *PAPSS2-PTEN* locus deletion in single cell-derived HAP1 clones transfected with the pBlueScript plasmid (17%, 1/6) or no plasmids (29%, 5/17). Since there was no significant difference in the frequency of the *PAPSS2-PTEN* locus deletion between HAP1 clones transfected with or without small size plasmid (**Fig. 5B**), we ruled out that the transfection of plasmids affected the *PAPSS2-PTEN* locus deletion.

To investigate when the *PAPSS2-PTEN* locus deletion appeared, we profiled consecutive passages of single cell-derived HAP1 cell clones. We found that the *PAPSS2-PTEN* locus deletion occurred predominantly in the first passage (96%, 45/47) and rarely in the second (2%, 1/47) or third (2%, 1/47) passage (**Fig. 5C**).

HAP1 is considered a near-haploid cell line but frequently transitions to the more stable diploid cell stage (42, 43). To investigate whether genome ploidy was linked to the loss of the *PAPSS2-PTEN* locus, we determined the degree of ploidy in 35 single cell-derived HAP1 cell clones with (n=18) or without (n=17) the *PAPSS2-PTEN* locus using a mixed population of haploid and diploid HAP1 cells as control. We found that the number of HAP1 cell clones residing in the haploid, haploid-to-diploid-transitioning or diploid stage was almost identical (**Fig. 5D, Supplementary Fig. S4A-C**), verifying that the ploidy status was neither causing nor affecting the deletion of the *PAPSS2-PTEN* locus.

Since PTEN and KLLN have been reported to inhibit cell proliferation, we tested whether the *PAPSS2-PTEN* locus deletion could provide HAP1 cells with a growth advantage. We did not observe a proliferative advantage in Δ*PAPSS2-PTEN* HAP1 cells without exposure to the CRISPR-Cas9 components or puromycin selection (**Supplementary Fig. S4A-B, Supplementary Fig. S4D-E**). However, when exposed to high levels of cellular toxicity, Δ*PAPSS2-PTEN* HAP1 cells were more likely to escaping cell cycle arrest, which may give those cells a growth advantage (**Fig. 5E, Supplementary Fig. S4C** and **S4E**).

In conclusion, the deletion of the *PAPSS2-PTEN* locus occurred at low frequency in HAP1 cells devoid of plasmid transfections or antibiotic selection. In contrast, the probability of losing the *PAPSS2-PTEN* locus significantly increased during the process of generating CRISPR-Cas9 cell clones and can make HAP1 cells more resilient to cellular stress.

### The *PAPSS2-PTEN* locus deletion in HAP1 cells was evident in human cancers

Since we found that up to 30% of the HAP1 cell clones showed a deletion of the *PAPSS2-PTEN* locus without stressors applied (**Fig. 5B**), and under consideration that HAP1 cells originated from chronic myelogenous leukemia, we further investigated the occurrence of this deletion across cancer types using patient data.

First, we inspected gene aberrations of the *PAPSS2, ATAD1, KLLN* and *PTEN* loci in 26 cancer types. Overall, we found frequently occurring deep deletions of and mutations in the *PAPSS2* and *PTEN* gene as well as deep deletions of the *ATAD1* and *KLLN* gene (Materials and Methods). All four genes were deleted in 73% (19/26) of the assessed cancer types (**Fig. 6A**). The frequency of genomic alterations of these loci was relatively low in leukemia, but deep deletions were still highly prevalent. Alterations in cancer genomes are usually large local events, driven by tumor suppressor genes or oncogenes (44–46). When cancer cells lose tumor suppressor genes, the nearby genes can be collaterally deleted (44–46). We found that the *PTEN* gene locus was lost in 466 patients, many of which had also acquired deep deletions of the *KLLN* (67%), *ATAD1* (59%) and *PAPSS2* (43%) gene locus (**Fig. 6B**). Remarkably, these accumulative gene deletions reflected the gene order in the linear genome, suggesting that *PTEN* is the primary deletion event, and *KLLN, ATAD1* and *PAPSS2* are collaterally deleted (**Fig. 6B**).

**Fig 6.**
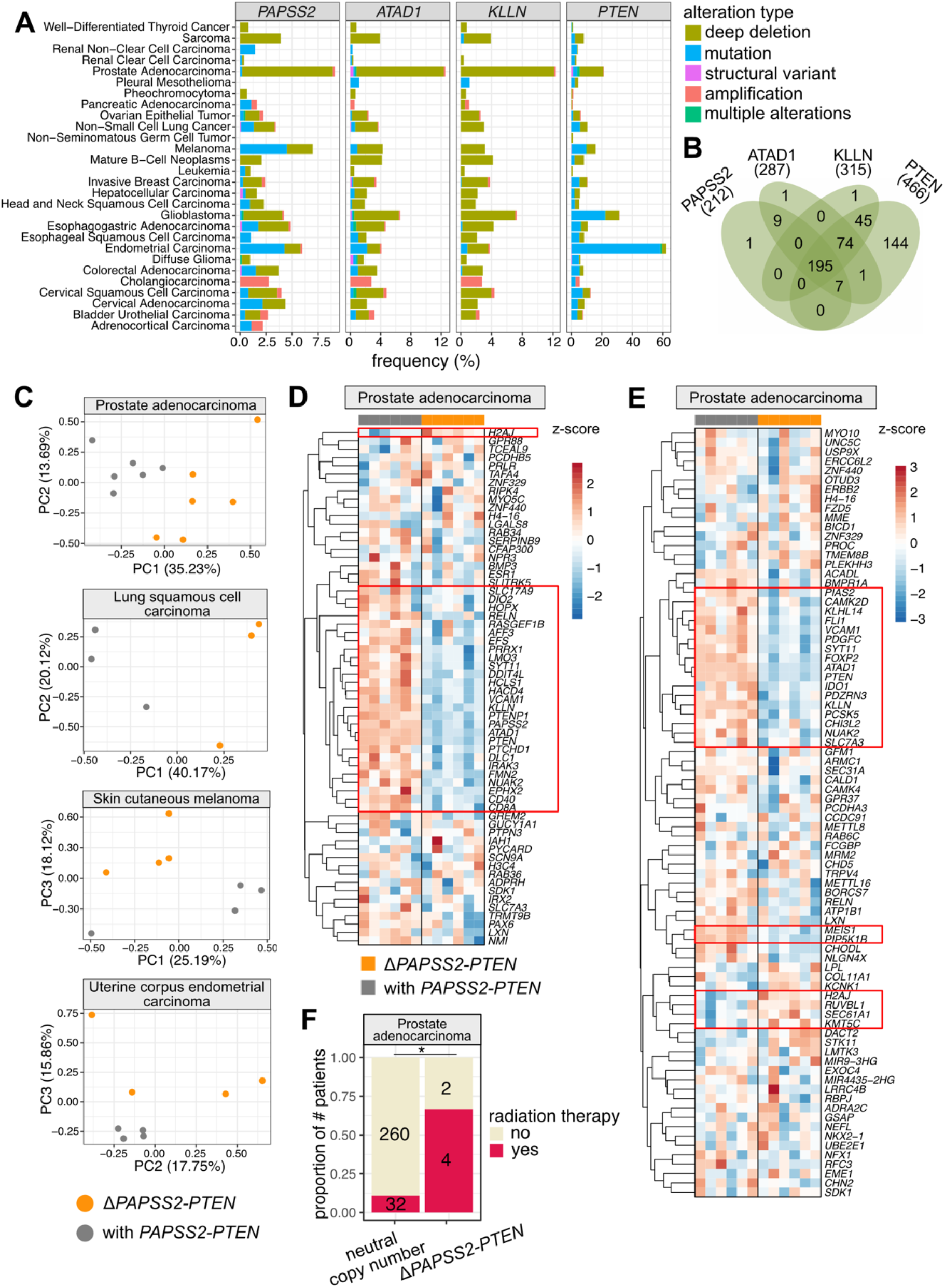
The deleted *PAPSS2-PTEN* locus occurred frequently in cancer patients. (**A**) Bar plots indicate the frequencies of genomic alterations occurring at the *PAPSS2, ATAD1, KLLN* and *PTEN* gene loci across 26 human cancer types. The data were retrieved from cBioPortal for Cancer Genomics (27). (**B**) Four-way Venn diagram intersects the numbers of patient samples carrying deep deletion at the *PAPSS2, ATAD1, KLLN* or *PTEN* gene loci. (**C**) PCA distinguishes patients with (grey) and without (orange) the *PAPSS2-PTEN* gene locus in prostate adenocarcinoma, lung squamous cell carcinoma, skin cutaneous melanoma and uterine corpus endometrial carcinoma. The proportion of variance explained by each PC is indicated in parenthesis. (**D-E**) Heatmaps show the expression levels of genes in prostate adenocarcinoma patients identified as DE genes in HAP1. The data were log_10_ transformed. Red: z-score > 0, blue: z-score <0. The (**D**) top 100 downregulated genes ordered by FC and (**E**) genes with H3K27 DAc sites located within 5 kb are shown. The patient samples on the top of each heatmap are colored by groups (orange: Δ*PAPSS2-PTEN*, grey: with *PAPSS2-PTEN*). Genes with consistent expression changes across the biological replicates are highlighted by red squares. (**F**) Stacked bar plot shows the fraction of prostate adenocarcinoma patients with (neutral copy number) and without the *PAPSS2-PTEN* gene locus upon radiation therapy. Statistics: Fisher’s exact test. Significance codes: *0.01<p<0.05.

Second, we investigated whether the collateral deletion at the *PTEN* locus is accompanied by transcriptional changes. For each cancer type, we searched for matching gene expression and copy number datasets of at least three patients carrying homozygous deletions of these four genes (*PAPSS2, ATAD1, KLLN* and *PTEN*) without considering other factors, such as tumor grade, age, or gender of the patients. Of the 23 cancer types, we found four cancer types (prostate adenocarcinoma, lung squamous cell carcinoma, skin cutaneous melanoma and uterine corpus endometrial carcinoma) with corresponding patient datasets. Based on gene expression, our PCA separated patient samples according to the occurrence of the *PAPSS2-PTEN* locus deletion in the first three PCs (**Fig. 6C**), which was striking given that other factors would usually distinguish individual patient samples.

Third, we examined to which extent the altered gene expression pattern identified in HAP1 cells was reflected in cancer patients. Genes downregulated in the Δ*PAPSS2-PTEN* HAP1 cells showed also reduced gene expression in Δ*PAPSS2-PTEN* patient samples (**Fig. 6D, Supplementary Fig. S5A**). For example, 13% to 45% of the top 100 downregulated genes in HAP1 were also deregulated in prostate adenocarcinoma, lung squamous cell carcinoma and skin cutaneous melanoma when comparing patients with and without the *PAPSS2*-*PTEN* locus deletion (**Supplementary Table S13**). Similarly, some of the DE genes which were located within 5 kb of a H3K27 deacetylated site in HAP1 cells also exhibited differences in gene expression in patient samples with and without the *PAPSS2-PTEN* locus deletion (**Fig. 6E, Supplementary Fig. S5B, Supplementary Table S13**).

Lastly, we observed that the frequency of the *PAPSS2-PTEN* locus deletion significantly increased in prostate cancer patients with radiation therapy (**Fig. 6F**). This finding agreed with our observation that the frequency of the collateral deletion dramatically increased when HAP1 cells were exposed to puromycin in the absence of a resistance gene (**Fig. 5B**).

In summary, the deletion of the *PAPSS2-PTEN* locus resulted in similar gene expression changes across different cancer types and may provide cancer cells with a selective advantage when exposed to external stressors, such as upon cancer treatments.

## DISCUSSION

Frequent genomic alterations, including the deletion at 10q23, have been detected in cancer cell lines (47–49). Studying cancer cell lines with known and well-characterized genomic deletions allows to extract essential clinical and pharmacological relevant information (47–50). However, the 10q23 (Δ*PAPSS2*-*PTEN*) deletion remained so far unnoticed in CRISPR-Cas9-modified HAP1 cell clones. Furthermore, the deletion occurs only in a fraction of HAP1 cell clones resulting in a heterogenous pool of HAP1 cell clones with and without the *PAPSS2*-*PTEN* locus skewing biological observations. Therefore, selecting correctly CRISPR-Cas9-modified cell clones with stable genotypes is critical for reliable interpretations of disease phenotypes.

In addition, it is important to consider that many treatments used in cell-based assays are genotoxic and trigger genome instability. For example, antibiotic treatment with puromycin is generally stressful for mammalian cells. As a potent inhibitor of protein synthesis, puromycin will inhibit cell growth in a dose- and exposure time-dependent manner and especially effects cells without the puromycin resistance gene. Accordingly, we observed that the absence of the puromycin resistance gene increased the frequencies of the *PAPSS2*-*PTEN*. However, we also need to emphasize that up to 30% of all single cell-derived HAP1 cell clones carry the *PAPSS2*-*PTEN* deletion even without puromycin selection. Thus, despite the px459 mediated puromycin resistance in our CRISPR-Cas9-modified HAP1 cell clones, the increased occurrences of the *PAPSS2*-*PTEN* deletion in the CRISPR-Cas9-modified cells were likely the consequence of gRNA-independent Cas9 protein-mediated toxicity (51–53). Furthermore, Cas9-induced DSB can activate DNA damage checkpoints (54). Since *PTEN* and *KLLN* encode proteins regulating the G1-S transition (55) and cell cycle arrest coupled to apoptosis (37), respectively, Δ*PAPSS2-PTEN* cells could be faster released from cell cycle checkpoints. Accordingly, during the puromycin selection process, cells without the resistance gene will be exposed to genotoxic stress (56) and thus, the *PAPSS2-PTEN* locus deletion would allow cells to escape cell arrest and consequently survive.

Our findings in HAP1 cell lines underscored the clinical significance of the genomic deletion at 10q23.3 for cancer patient outcome. Previous studies reported the frequent loss of the *PAPSS2*-*PTEN* locus in prostate cancer and its association with prostate-specific antigen reoccurrence in patients. In addition to impairing the tumor suppressing roles of PTEN, the ablated metabolic functions of PAPSS2 have been linked to cancer reoccurrences, which emphasize the combined impact of collateral gene deletions in cancer cells (57, 58). Additive effects of deleted genes were reported in other cancer types and genomic loci as well. For example, glioma cells carrying a large deletion including the *ENO1* gene cannot survive when the paralogue ENO2 was inhibited. Conversely, the depletion of ENO2 only marginally limited cell proliferation when *ENO1* remained intact (46). The vulnerability created by collateral deletions in cancer cell provides opportunities for specific and efficient cancer treatment options. Thus, the increase in cancer cell apoptosis upon collateral deletion of genes in the *PAPSS2*-*PTEN* locus can be exploited in treatment strategies through a targeted co-deletion of *ATAD1* and *PTEN* (59).

Although the precise mechanism that initiated the *PAPSS2-PTEN* locus deletion remains unclear, our findings indicated mechanistic similarities linked to genome fragility. Genome-wide screening identified hundreds of fragile sites leading to DNA breaks initiated upon cell treatment with DNA replication inhibitors (60, 61). Among them, the *PAPSS2-PTEN* locus resided in the rare fragile site FRA10A (10q23.3). Fragile sites usually form DNA structures that differ. to the classic B-DNA helix, such as R-loops, G-quadruplexes or stem loops, which can hinder DNA replication, leading to replication fork stalling and DNA breaks (62). Previously, it has been shown that *PTEN* exon 1 forms a highly stable secondary structure *in vitro* (63). In accordance, we identified potential G-quadruplex structures in exonic sequences of *PTEN, KLLN, ATAD1* and *CFL1P1* in human cells (**Supplementary Fig. S6A**) (33). Moreover, our detailed inspection of the sequence content revealed the possible formation of non-B-DNA structures by A-mononucleotide repeat sequences and GT-repeats at the DNA break boundaries of the deleted *PAPSS2-PTEN* locus (**Supplementary Fig. S6B**). We noticed (A)_37_ and (A)_30_ repeat stretches occurring downstream of *PTEN* gene body that can trigger slipped-strand DNA structures corresponding to single stranded DNA loops interspersed within double stranded DNA (62). A-mononucleotide repeat sequences can act as DNA unwinding elements (64, 65) initiating fork stalling and collapse that results in DNA breaks upon treatment with agents inhibiting DNA synthesis (64). We found (GT)_20_ repeats located in the first intron of *PAPSS2* that can result in left-handed Z-DNA and induce DSBs in mammalian cells (66). Altogether, we reasoned that the *PAPSS2-PTEN* locus is highly vulnerable upon exposure to DNA replication stress.

HAP1 cells were generated as a by-product of transfecting KBM-7 cells with Yamanaka factors to obtain iPSCs (67). The two Yamanaka factors c-MYC and KLF4 are proto-oncogenes. Oncogenes are known to generate replication stress (68). Moreover, concerns of the genome instability in iPSCs have been raised for years (69). Therefore, we speculate that the exogenous replication stress exerted when HAP1 cells had been generated made them more prone to DSB at the *PAPSS2-PTEN* locus when compared to other cell lines. Similarly, DNA replication stress increases considerably during tumor initiation and progression and might explain the high occurrence of *PAPSS2-PTEN* locus deletion in cancer patients.

We found that cancer cells carrying the *PAPSS2-PTEN* locus deletion exhibited significant changes on multiple levels ranging from global changes in the chromatin environment to cell behavior. This can confound our understandings of cancer cells, especially since HAP1 cells are commonly used in large-scale CRISPR-Cas9-based genetic screens or targeted functional studies of cancer phenotype-associated genes (9). Thus, awareness of the dramatic genomic alterations is crucial for CRISPR-Cas9 applications. Furthermore, the molecular characteristics of the *PAPSS2-PTEN* locus deletion that we identified in HAP1 cells can be linked to the commonly observed collateral deletion of these genes in cancer patients. Therefore, our observation underscored the necessity of rigorous HAP1 cell clone validation experiments when applying CRISPR-Cas9-mediated gene editing experiments, and also highlighted the clinical relevance when investigating the impact of collateral gene deletions in cancer patients and the responsiveness to cancer treatments.

## MATERIAL & METHODS

### Cell culture

HAP1 cells were purchased from Horizon Discovery with a certified genotype, mycoplasma-free tested and grown in Iscove Modified Dulbecco Medium (IMDM, HyClone™) supplemented with 10% Fetal Bovine Serum (FBS, HyClone™) and 1% Penicillin-Streptomycin (Sigma-Aldrich). Cells were cultured in T75 flasks at 37°C and 5% CO_2_. HAP1 cells were expanded by splitting 1/10 every 2 days or when reaching 50% to 60% confluency. Upon splitting, the medium was aspirated, cells were washed with phosphate buffered saline (PBS, Sigma-Aldrich) and then detached with 2 mL of a trypsin-EDTA solution (Sigma-Aldrich). Trypsin was subsequently inactivated by adding a minimum of 3-fold surplus of medium.

### Plasmid construction, transfection and generation of single cell-derived clones

gRNAs were designed and the targeting potential assessed (https://zlab.bio/guide-design-resources). Each gRNA was individually cloned into the two BbsI restriction sites pSpCas9(BB)-2A-Puro (px459). For generating non-targeting control clones, px459 without any gRNA sequence was transfected. To enhance transfection efficiency, pBlueScript was co-transfected with CRISPR-Cas9 plasmid at a 1:1 ratio (10). One day before transfection, about 160,000 HAP1 cells were plated in each well of a 6-well plate, and on the next day transfected using TurboFectin 8.0 (OriGene) according to the manufacturer’s instructions. The cells were incubated together with the plasmid-TurboFectin mixture for 24 h and then selected by adding fresh cell culture medium with 2 μg/mL puromycin for 48 h. Afterwards, cells recovered in the complete medium without puromycin. After recovery, about 100-500 cells were seeded into 10 cm dishes to form single cell-derived clones. Those clones were hand-picked under the microscope and expanded.

### Genome extraction

Cells were lysed in 400 μL lysis buffer (0.5% SDS, 0.1M NaCl, 0.05M EDTA, 0.01M Tris-HCl, 200 μg/mL proteinase K). After overnight incubation at 55ºC, 200 μL of 5M NaCl were added, and the sample was vortexed and incubated on ice for 10 min. After centrifugation (15,000×*g*, 4ºC, 10 min), 400 μL of the supernatant was transferred to a new tube and mixed with 800 μL of 100% ethanol. The samples were incubated on ice for at least 10 min. Genomic DNA was pelleted by centrifugation (18,000×*g*, 4ºC, 15 min,) washed once with 70% ethanol and resuspended in nuclease-free water.

### PCR

PCR primers were designed using NCBI Primer-BLAST with default parameters. PCR was performed with Taq polymerase (New England Biolabs) according to the manufacturer’s instructions. Briefly, the 25 μL PCR reaction was used (1× standard Taq reaction buffer, 200 μM dNTPs, 0.2 μM primers (**Supplementary Table S1**), 1U Taq DNA polymerase and 100-1000 ng of genomic DNA). PCR was completed by an initial denaturation step (95°C for 5 min), 35 amplification cycles (95°C for 30 sec, 55-60°C for 30 sec or for at 68°C for 60 sec per 1 kb DNA), a final extension step (68°C for 5 min) and holding the reaction at 4°C in a thermocycler (Applied Biosystems) with a preheated lid (105°C). The products were assessed by 1.2% agarose gel electrophoresis.

### Quantitative PCR (qPCR)

Genomic template DNA was incubated at 37°C to homogenously resuspend. About 10-100 ng genomic DNA with 2.5 μM primers targeting the promoter region of *PTEN* or *ATAD1* and a genomic region on Chr 12 as an internal control (**Supplementary Table S1**) and PowerUp™ SYBR™ Green Master Mix (Applied Biosystems™) were mixed. qPCR was performed using an initial denaturation step (50°C for 2 min, 95°C for 2 min), 40 amplification cycles (95°C for 15 sec, 60°C for 1 min) and a step for obtaining the melting curve (95°C for 15 sec, 60°C for 1 min and ramp rate 1.6°C/sec, 95°C for 15 sec and ramp rate 0.075°C/sec) in a QuantStudio5 Real-Time PCR System (Thermo Fisher Scientific).

### Ploidy and cell cycle assessment

Ploidy level and cell cycle were assessed by flow cytometry (BD LSR II SORP with BD FACSDiva™ software version 9.0, BD Biosciences) by gating live cells in FSC-A/SSC-A plot and singlets in FSC-H/FSC-A plot. DNA content was measured with a 561 nm laser. The results were further analyzed with the FlowJo software version 8.2 and the Dean-Jeff-Fox algorithm for cell cycle analysis and visualization. About 500,000 cells were collected and washed once in PBS. Cells were fixed by adding of 500 μL of ice-cold 70% ethanol drop-wise while vortexing, and subsequently stored at -20ºC until further processing. For the cell staining, fixed cells were pelleted and washed twice with PBS and resuspended in 300 μL hypotonic buffer (1 g/L sodium citrate buffer, 0.1% Triton-X 100) supplemented with 40 μg/mL propidium iodide (Sigma-Aldrich) and 100 μg/mL of RNase A (Thermo Fisher Scientific). A haploid and diploid cell population was used as gating reference.

### Cell proliferation assay

About 1000 cells were seeded per well in sextuplet in a 96-well plate, and the cell number was measured daily over 4 days (day 0, 1, 2, and 3). Upon measurement, the culture medium was aspirated, and cells were incubated by a mixture of 60 μL of IMDM and 10 μL of MTT (4 mg/mL Methylthiazolyldiphenyl-tetrazolium bromide, Sigma-Aldrich, in PBS) at 37ºC for 75 min. Next, the supernatant was replaced by 100 μL lysis buffer (90% isopropanol, 0.5% SDS, 0.04N HCl) and cells were incubated on a rocker at room temperature for 30 min. Lysed cells were resuspended and measured on a plate reader (Molecular Devices Spectramax i3x, 595 nm absorbance). Wells without any seeded cells were used as background control. To obtain the optical density (OD), background values were subtracted from the obtained signals per well with seeded cells.

### Chromatin immunoprecipitation followed by sequencing (ChIP-seq)

ChIP-seq experiments were performed as previously described (11, 12). Briefly, 15-20M cells were fixed (1% formaldehyde), lysed, sonicated (Covaris ME220, milliTUBE 1 ml AFA Fiber, with parameter: Setpoint Temperature 9°C, Peak Power 75, Duty Factor 15%, Cycles/Burst 1000, Duration: around 20 min) and then incubated with H3K4me3 (05-1339, Millipore) and H3K27ac (ab4729, abcam) antibodies. After chromatin immunoprecipitation, sequencing libraries were generated (Takara SMARTer^®^ ThruPLEX^®^ DNA-seq Kit following the manufacture’s protocol), library quality and size distribution were assessed (Agilent Bioanalyzer, High Sensitivity DNA chips) and quantified (KAPA SYBR^®^ FAST qPCR kit, Roche). Sequencing was performed with the NextSeq 500/550 High Output v2 kit for 75 cycles (Illumina), single-end Nextseq500 (Illumina).

### ChIP-seq data analysis

The qualities of reads were assessed by FastQC (13). Reads were aligned to the human reference genome (hg38) using BWA (14). After alignment, PCR duplicates and reads mapping to the ENCODE exclusion list (https://sites.google.com/site/anshulkundaje/projects/blacklists) were removed using SAMtools (15) and NGSUtils (16). The bam files were indexed and sorted by SAMtools. MACS2 was used for peak calling (17). Subsequently, differential enrichment analysis was performed with DiffiBind (18). Significantly DAc peaks (FDR ≤0.05) were identified. The distances between individual DAc peaks to the nearest DE genes and individual DE genes to the nearest DAc peaks were calculated using BEDTools (19). For visualization, bedgraph files were generated using deepTools (20). Reads with a mapping quality below 20 were removed with SAMtools, and bedgraph files were visualized with IGV (21).

### RNA extraction and DNase-treatment

Cells were harvested at 50% to 60% confluency. Between 1,000,000 to 5,000,000 cells were pelleted and lysed in 700 μL Qiazol (QIAGEN). Afterwards, 140 μL chloroform were added. The mixture was shaken for 30 sec and incubated for 2.5 min at room temperature, followed by centrifugation (9,000×*g*, 4°C, 5 min) to achieve phase separation. The upper aqueous phase was carefully transferred to a new tube, and 1 volume isopropanol was added. The tubes were inverted 5 times to mix thoroughly, followed by incubation (room temperature, 10 min). The RNA was pelleted by centrifugation (9,000×*g*, 4°C, 10 min) and washed once with ice-cold 70 % ethanol. The RNA was resuspended in 30-50 μL nuclease-free water. RNA concentration and purity were determined (NanoDrop™ 2000c). To remove genomic DNA, 10 μg RNA were mixed with 5 μL 10×TurboDNase buffer (Invitrogen), 1 μL TurboDNase (Invitrogen), 1 μL RNase Inhibitor (RiboLock, Invitrogen), and water was added to a total volume of 50 μL. The samples were incubated (37°C, 30 min) and purified with the Zymo RNA Clean & Concentrator Kit (Zymo Research). The concentration of purified RNA was determined (Qubit™ RNA HS Assay Kit), and the integrity was assessed (Agilent Bioanalyzer, RNA 6000 Nano kit).

### RNA-seq

The RNA was first enriched for molecules with PolyA tails using NEBNext^®^ Poly(A) mRNA Magnetic Isolation Module. The RNA library preparation was performed with NEBNext^®^ Ultra™ II Directional RNA Library Prep Kit for Illumina according to the manufacturer’s instruction. Library quality was determined (Agilent Bioanalyzer, High Sensitivity DNA chips) and quantified (KAPA SYBR^®^ FAST qPCR kit, Roche). Sequencing was performed with the NextSeq 500/550 High Output v2 kit for 150 cycles (Illumina) paired-end on Nextseq500 (Illumina).

### RNA-seq data analysis

Sequencing read qualities were assessed with FastQC. Adaptor sequences and low-quality reads were trimmed using Trimmomatic (22). Sequencing reads that can be aligned (HiSAT2) (23) to annotated ribosomal RNA genes were discarded. Subsequently, the filtered reads were aligned to the human genome hg38 using HiSAT2. Using sorted and indexed bam files, the number of aligned reads were counted for each annotated transcripts (featurecount in subread package) (24). For visualization, bedgraph files were generated using deepTools with soft-clipped reads removed by SAMtools. The raw count tables were used to identify differentially expression genes (FDR ≤ 0.05) with DESeq2 (25). The over-representative enrichment of Gene ontology (GO) term and KEGG pathway analyses were performed and visualized using ClusterProfiler (26).

### Additional data sources

Gene expression data across different cell lines (**Fig. 1C**) were retrieved from the Human Protein Atlas (v. 21.0) (https://www.proteinatlas.org/about/download) under the section RNA HPA cell line gene data (28). Published Hi-C data on HAP1 cells (**Fig. 1D**) were visualized using 3D Genome browser (29, 30). Published ChIA-PET data for multiple human cell lines (**Fig. S1C**) were visualized through WashU Epigenome Browser (31). Interaction networks (**Fig. S2A**) were generated with STRING database V11.5 (32). G-quadruplex site prediction and structure mapping using CUT&Tag in HEK293T cells (**Fig. S6A**) were performed by Jing *et al* (33).

## List of abbreviations

Δ: deletion
3D: three-dimension
Chr: Chromosome
ctrl: control clones
BP: biological process
Cas9: CRISPR-associated protein 9
ChIP-seq: chromatin immunoprecipitation followed by sequencing
CRISPR: clustered regularly interspaced short palindromic repeats
DE: differentially expressed
DAc: differentially acetylated
DSB: double-strand break
FC: fold-change
FDR: false discovery rate
gRNA: guide RNA
H3K4me3: histone 3 lysine 4 trimethylation
H3K27ac: histone 3 lysine 27 acetylation
iPSCs: induced pluripotent stem cells
GO: gene ontology
MF: molecular function
PCA: Principal component analysis
Pol II: polymerase II
tRNA: transfer RNA
TAD: topologically associating domain
TSS: transcription start site

## ACKNOWLEDGEMENTS

We would like to thank the groups of Claudia Kutter, Marc Friedländer, and Vicent Pelechano group, as well as Ian Mills, Laura Baranello, Marcel Van Vugt, Philip Yuk Kwong Yung, Jing Lyu and Bruno Urién for the critical discussion and feedback. We thank the Swedish Bioinformatics Advisory Program and Erik Fasterius at the National Bioinformatics Infrastructure Sweden for advice. Computations were enabled by resources in project SNIC 2017/7-261, SNIC 2020/16-223, SNIC 2020/15-292, SNIC 2021/22-899, SNIC 2021/23-652, SNIC 2022/22-1063 and SNIC 2022/23-546 provided by the Swedish National Infrastructure for Computing at UPPMAX.

