## Supplemental Figures for "CRISPR-Cas9-mediated genome engineering exaggerates genomic deletion at 10q23.31 including the *PTEN* gene locus mimicking cancer profiles"

### SUPPLEMENATRY FIGURES & FIGURE LEGENDS

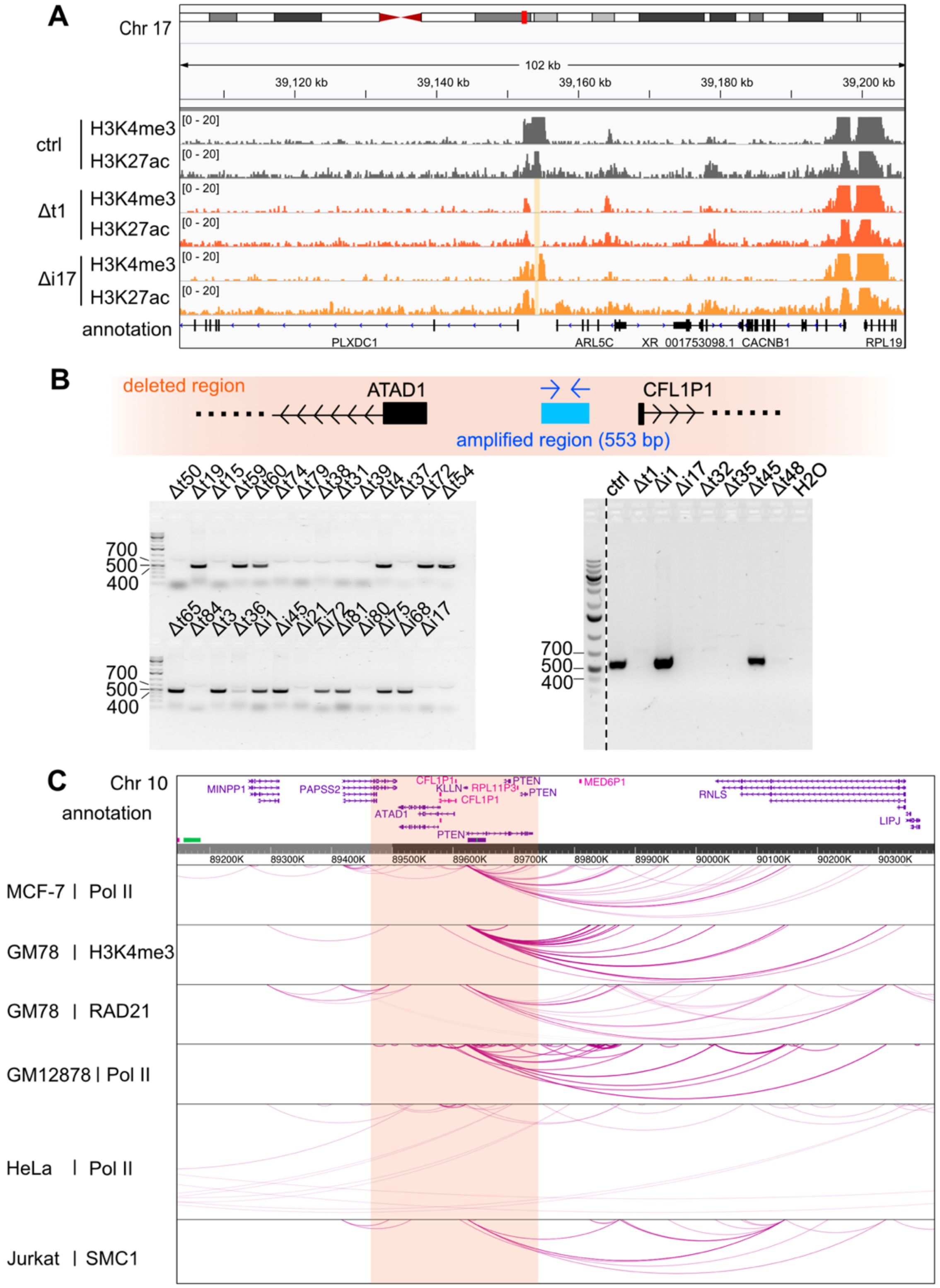


**Fig S1. The loss of *PAPSS2-PTEN* locus was confirmed in CRISPR-Cas9-modified HAP1 single cell-derived clones**

(**A**) The hg38 genome browser (zoom-out view of Fig. 1A) illustrates normalized H3K4me3 and H3K27ac ChIP-seq reads in control (ctrl, dark grey), Δt1 (orange) and Δi17 (yellow) HAP1 cell clones at the corresponding targeted loci on Chr 17 (beige box). (**B**) Schematic illustration shows annealed region of PCR primers and amplicon length designed to validate the presence (553 bp PCR product) or absence (no PCR product) of the *PAPSS2-PTEN* locus (*top*) in HAP1 Δt and Δi cell clones. Agarose gel confirms the size of the obtained PCR products (*bottom*). (**C**) The hg38 genome browser displays long-range interaction loops centered on the *PAPSS2-PTEN* locus from publicly available Pol II, RAD21, H3K4me3 and SMC1 ChIA-PET data in multiple human cell lines.

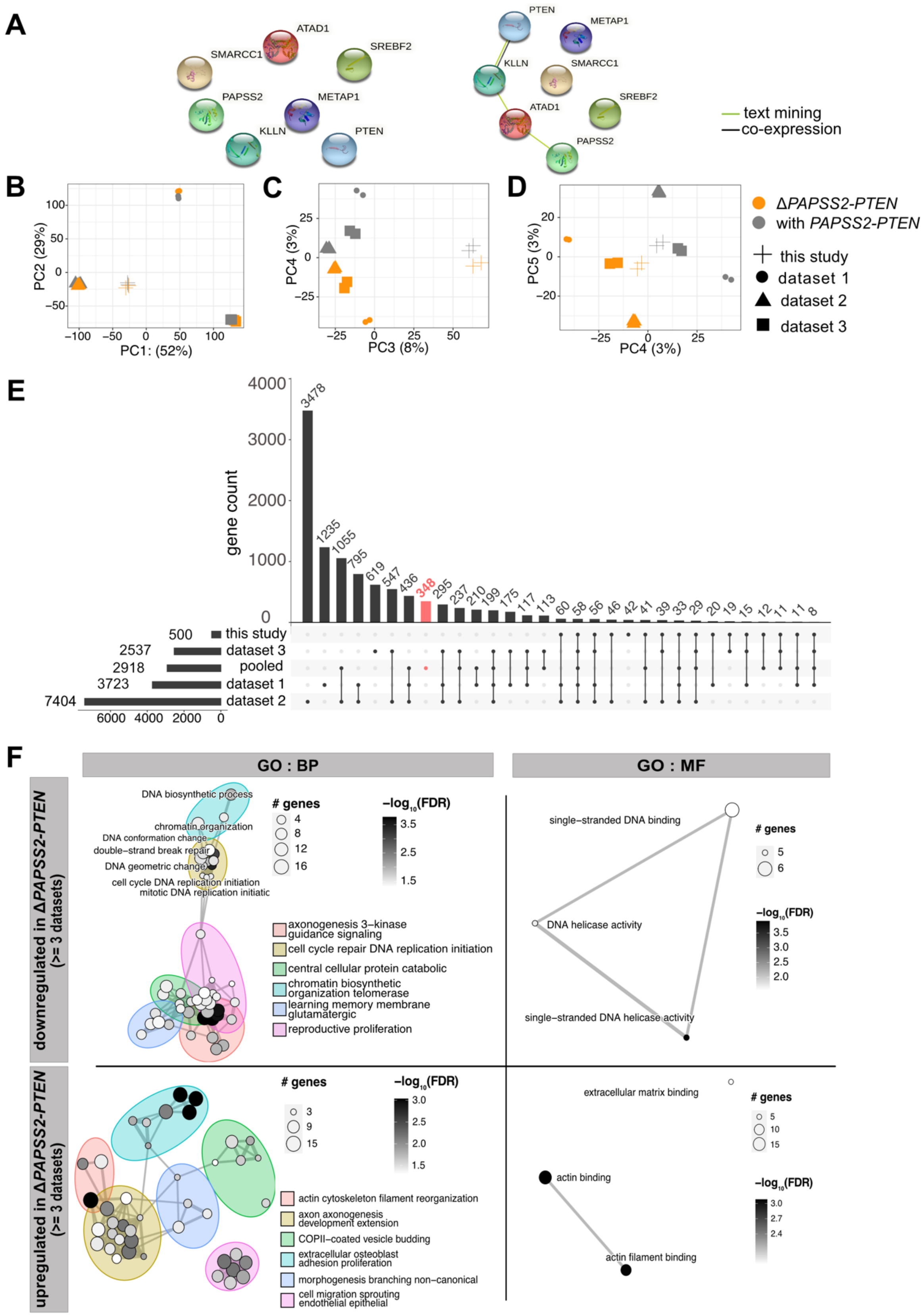


**Fig S2. Transcriptomic profiles were significantly altered in Δ*PAPSS2-PTEN* HAP1 cell clones**
(**A**) Interaction networks display connections between the four protein-coding genes located in the *PAPSS2-PTEN* locus and the CRISPR-Cas9-medidated target gene deletions of the published RNA-seq datasets (Fig. 2A). The minimum required interaction scores were set to 0.7 (high confidence, *left*) and 0.4 (medium confidence, *right*). (**B-D**) Factorial maps of the PCA for global gene expression without batch effect correction separate samples according to (**B**) the origin of samples (geometric symbols) and (**C-D**) absence (orange) or presence (grey) of the *PAPSS2-PTEN* locus in HAP1 CRISPR-Cas9 deletion clones. The proportion of variance explained by each PC is indicated in parenthesis. (**E**) UpSet plot intersects the number of differentially expressed (DE) genes (y-axis) across the four datasets (this study, published dataset

1 to 3) and all four datasets combined after batch effect correction (pooled). The red bar highlights the number of DE genes in the pooled datasets that do not overlap with DE genes identified in any of the datasets (this study and dataset 1, 2 and 3). (**F**) Enrichment maps illustrate all significantly enriched biological process (BP) and molecular function (MF) GO terms for down- and upregulated DE genes that were shared by at least three datasets. Each node represents one enriched GO term. The nodes are colored based on FDR-adjusted *p* values. The sizes of the nodes are determined by the numbers of DE genes contributing to the enriched GO term.

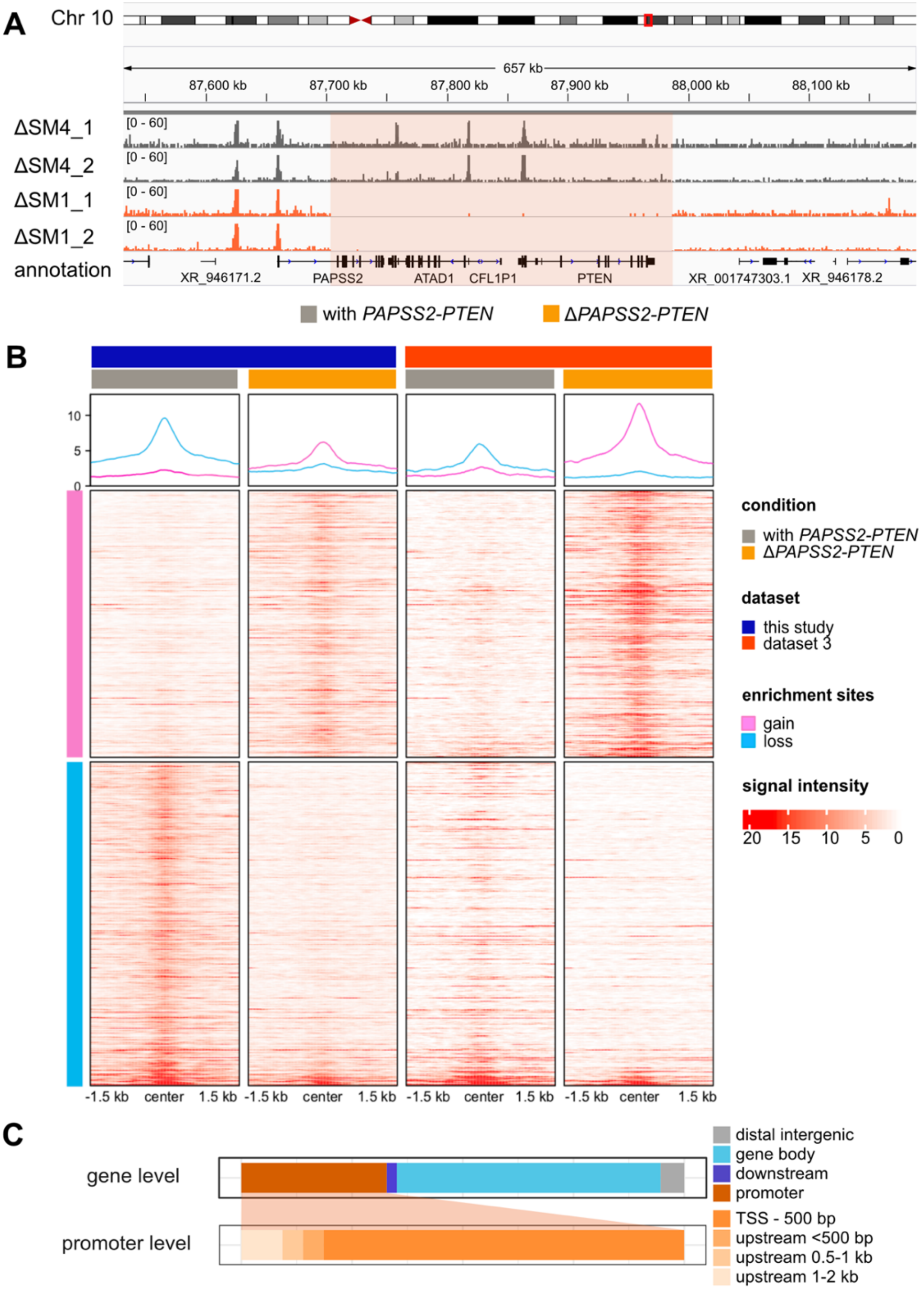


**Fig S3. Genome-wide differential acetylation of H3K27 occurred in Δ*PAPSS2-PTEN* HAP1 cell clones** (**A**) The hg38 genome browser displays normalized H3K27ac ChIP-seq reads around the *PAPSS2-PTEN* locus (red box) in two replicates of HAP1 cell clones with (ΔSM4, dark grey) or without (ΔSM1, orange) the *PAPSS2-PTEN* locus. (**B**) Heatmaps and aggregated plots show H3K27ac enrichment (red: H3K27ac bound to DNA and white: no binding identified) within 1.5 kb of the peak summit over all identified differentially acetylated regions (one per line, pink: increased and blue: reduced H3K27ac) in HAP1 cell clones with (grey) and without (orange) the *PAPSS2-PTEN* locus that were generated in this study (dark blue) and in dataset 3 (red). Two replicates are merged into one plot. (**C**) Stacked bars demonstrate proportional frequencies of a subset of differentially acetylated H3K27ac peaks near differentially expressed genes (DAcDE ≤ 5kb). Subcategories for promoter features are further divided.

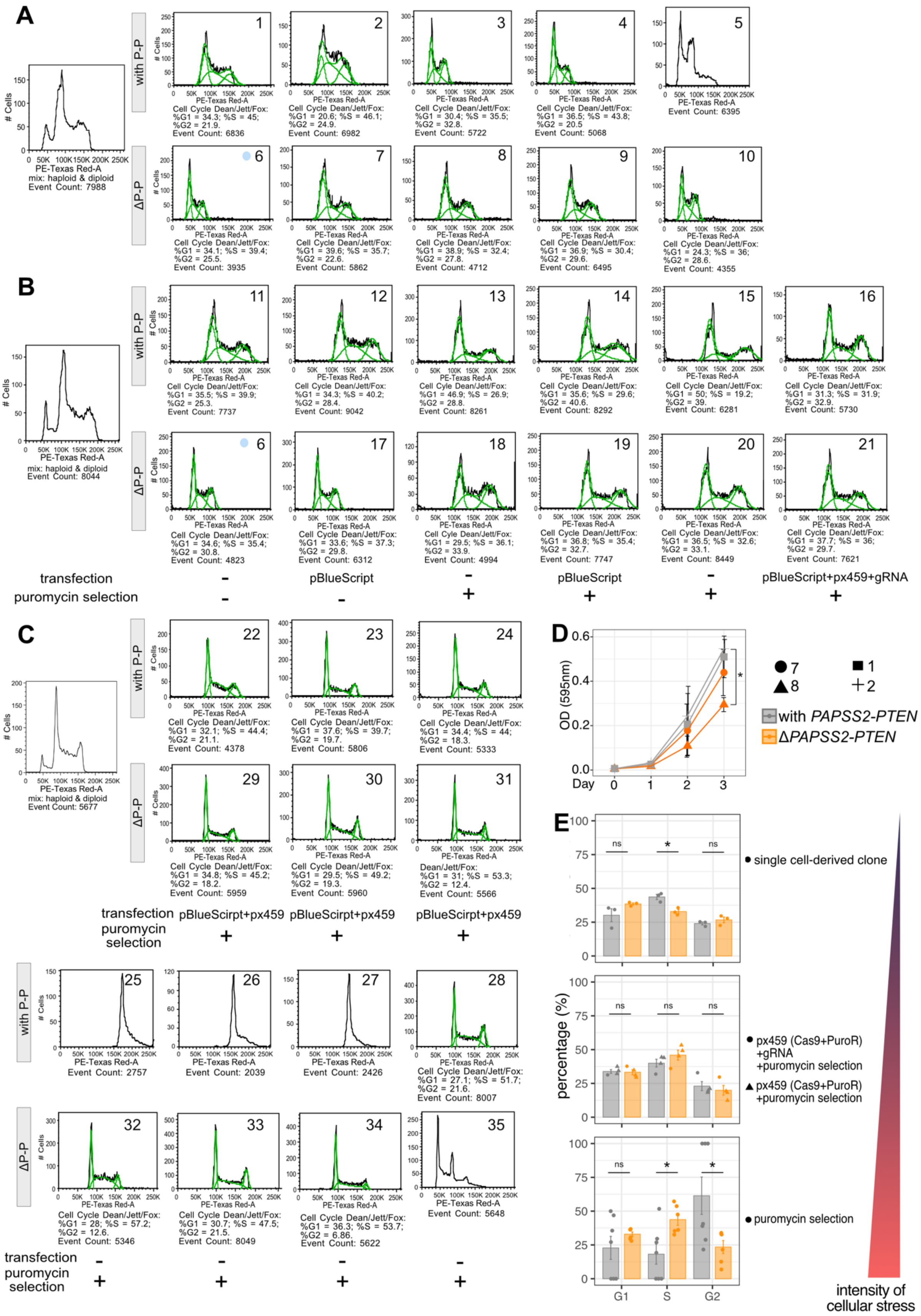


**Fig S4. The genomic deletion of the *PAPSS2-PTEN* locus changed the cell cycle**
(**A-C**) Flow cytometry plots (gating for alive singlets) display cellular ploidy of individual cells of the HAP1 single cell-derived clones with (P-P) or without (ΔP-P) the *PAPSS2-PTEN* locus. All HAP1 cell clones displayed under (**A**) underwent neither plasmid transfection nor puromycin selection, while the panel of HAP1 cell clones obtained after (**B-C**) plasmid transfection and puromycin selection or only plasmid transfection or only puromycin selection. The experimental settings are labelled below the plots. A mixed population of haploid and diploid HAP1 cells was used as the reference to determine genome ploidy (*left*).

Green lines indicate the event count and the number below each plot corresponds to the percentage of cells for each cell cycle phase (calculated by Dean/Jett/Fox model). Clone 6 was analyzed twice (blue circle in A and B). (**D**) Line graph shows the relative number of metabolically active cells (assessed by optical density, OD) measured over three days after seeding by MTT assay (n=2, mean +/- SD). Geometric symbols represent the different experimental settings. Statistics: two-tailed t-test. Significance codes: *0.01<*p*<0.05, insignificance is unlabeled. (**E**) The bar plots display the percentages of diploid HAP1 single cell-derived clones (same as in Fig S4. A-C) during the cell cycle phases. HAP1 cell clones are separated according to cellular exposure (*left:* mild, *right*: high cellular stress). Individual biological replicate (clone) is displayed. Statistics: two-tailed t-test. Significance codes: *0.01<*p*<0.05; ns, not significant.

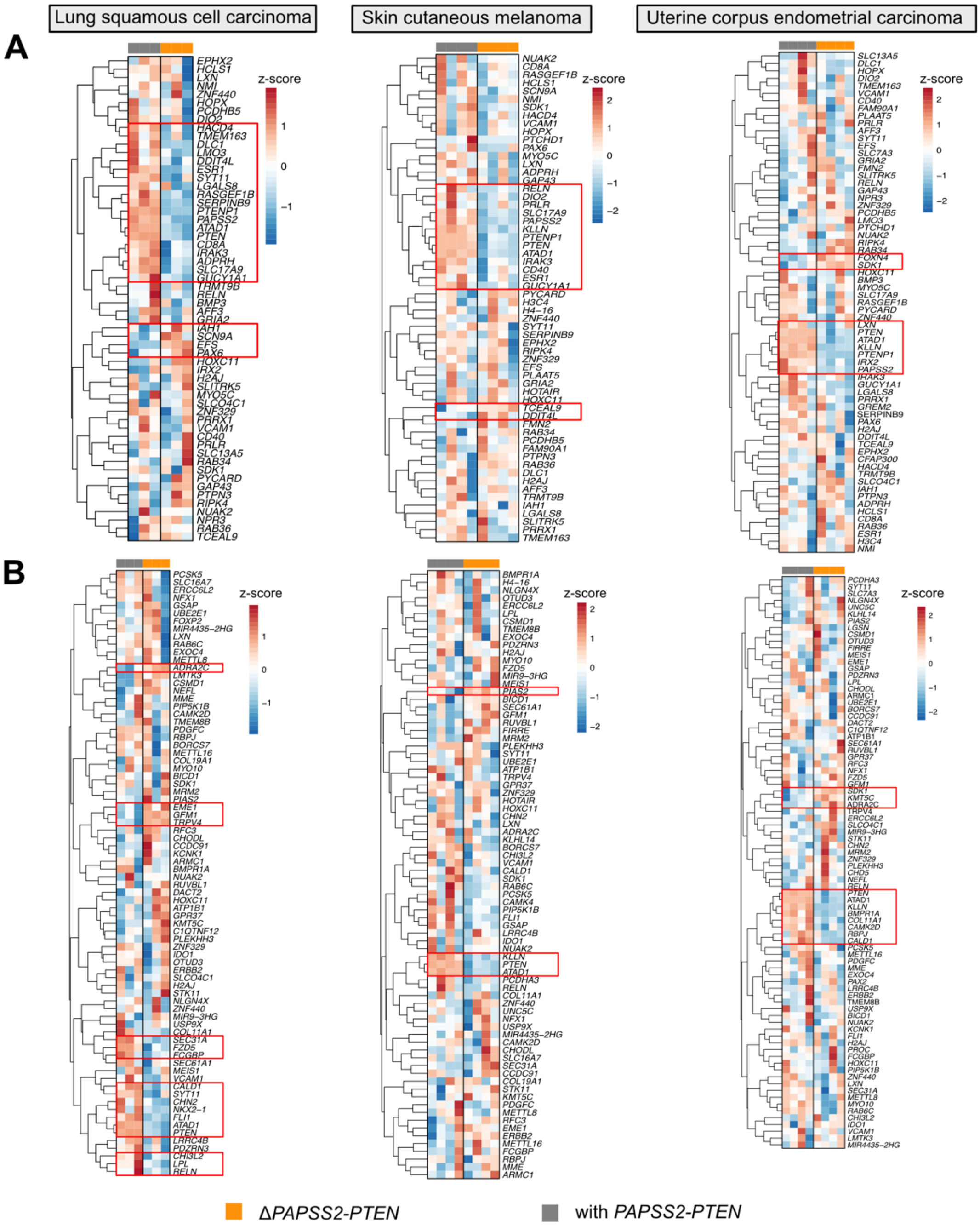


**Fig S5. Deletion of the *PAPSS2-PTEN* locus resulted in gene expression changed in cancer patients** (**A-B**) Heatmaps show the expression levels of genes in lung squamous cell carcinoma (*left*), skin cutaneous melanoma (*middle*) and uterine corpus endometrial carcinoma (*right*) patients identified as differentially expressed genes in HAP1 cells. The data were $\log_{10}$ transformed. Red: z-score > 0, blue: z-score <0. The (**A**) top 100 downregulated genes ordered by FC and (**B**) genes with differentially acetylated H3K27 sites located within 5 kb are shown. The patient samples on the top of each heatmap are colored by groups (grey: with *PAPSS2-PTEN*, orange: Δ*PAPSS2-PTEN*). Genes with common gene expression changes across the biological replicates are highlighted by red squares.

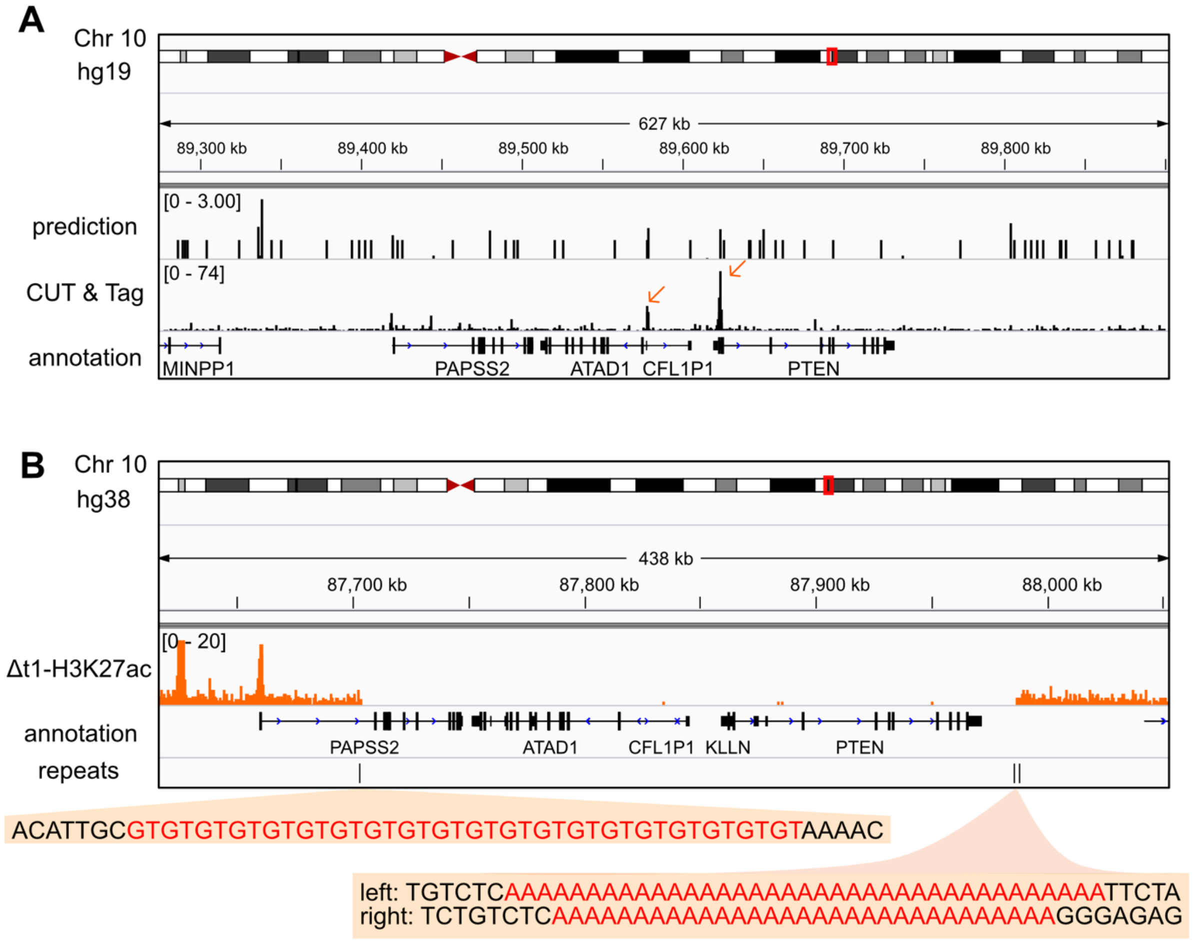


**Fig S6. Non-B DNA structures can form within the *PAPSS2-PTEN* locus and surrounding the DNA break sites**

(**A**) The hg19 genome browser displays genomic sites that are predicted to form G-quadruplex structure sites (upper track) and G-quadruplex CUT&Tag reads mapping to the *PAPSS2-PTEN* locus in HEK293T cells (lower track). The peaks located in the deleted region are highlighted (orange arrows). (**B**) The hg38 genome browser demonstrate H3K27ac ChIP-seq reads (orange) at the *PAPSS2-PTEN* locus in HAP1 Δt1 clone. The location of the repeats found around the DNA break site boundaries are shown (black vertical lines). Repeat sequence content (red) are highlighted (orange boxes) below the corresponding vertical lines.
